# Infralimbic projections to the basal forebrain mediate extinction learning

**DOI:** 10.1101/2024.06.19.599805

**Authors:** Carolina Fernandes-Henriques, Yuval Guetta, Mia Sclar, Rebecca Zhang, Yuka Miura, Allyson K. Friedman, Ekaterina Likhtik

## Abstract

Fear extinction learning and retrieval are critical for decreasing fear responses to a stimulus that no longer poses a threat. While it is known that the infralimbic region (IL) of the medial prefrontal cortex mediates retrieval of an extinction memory through projections to the basolateral amygdala (BLA), the contribution of the IL to extinction learning is not well-understood. Given the strong projection from the IL to the basal forebrain (BF), a center of attentional processing, we investigated whether this pathway participates in extinction, and compared it to the IL-BLA pathway. Using retrograde tracing, we first demonstrate that projections from the IL to the BF originate from superficial (L2/3) and deep cortical layers (L5), and that they are denser than IL projections to the BLA. Next, combining retrograde tracing with labeling of the immediate early gene cFos, we show increased activity of the L5 IL-BF pathway during extinction learning and increased activity of the L2/3 IL-BLA pathway during extinction retrieval. Our *in vitro* recordings demonstrate that neurons in the IL-BF pathway become more excitable towards the end of extinction learning, but less excitable during extinction retrieval. Finally, using optogenetics we show that inactivation of the IL-BF pathway impairs extinction learning, leaving retrieval intact. We propose that the IL acts as a switchboard operator during extinction, with increased L5 IL-BF communication during learning and increased L2/3 IL-BLA communication during retrieval. Anxiety and stress-related changes in IL physiology could affect one or multiple lines of communication, impairing different aspects of extinction.

**Significance Statement:** Extinction of conditioned fear is a widely used behavioral approach to diminish fear, with projections from infralimbic prefrontal cortex to the amygdala known for mediating extinction memory retrieval. However, less is known about infralimbic pathways involved in extinction learning. We use neuroanatomical tracing, behavior, slice recordings, and circuit manipulation to show that infralimbic output to the basal forebrain, an attention processing center, is denser than to the amygdala, and is active during extinction learning rather than retrieval. Neurons in the infralimbic-basal forebrain pathway become more excitable as extinction learning progresses, and then less excitable during extinction retrieval. Moreover, inhibiting this pathway impairs extinction learning. Our findings identify different lines of communication the infralimbic cortex uses for extinction learning and retrieval.

## Introduction

The ventral portion of the medial prefrontal cortex (mPFC), known as the infralimbic (IL) region in rodents, is critical for fear extinction (Milad and Quirk 2002, Phelps et al. 2004, Milad et al. 2007, Sierra-Mercado et al. 2011). IL neurons fire during consolidation of extinction learning and during extinction retrieval (Milad and Quirk 2002, Burgos-Robles et al. 2007), inhibition of the IL impedes fear extinction memory consolidation and retrieval, whereas IL stimulation accelerates extinction acquisition and retrieval (Vidal-Gonzalez et al. 2006, Burgos-Robles et al. 2007, Laurent and Westbrook 2009, Sotres-Bayon et al. 2009, Do-Monte et al. 2015, Kim et al. 2016, Szeska et al. 2022). However, the pathways through which IL integrates various components of learning remain unknown, thereby complicating translatability to humans (Roberts and Clarke 2019). For example, overactivation of the IL in rodents and vmPFC in non-human primates drives movement-based behavioral and cardiovascular responses to threat (Halladay and Blair 2017, Alexander et al. 2020). Likewise, in rats, IL neural firing decreases during fear-associated defensive freezing (Giustino et al. 2016), suggesting that this region may bias behavior away from defensive freezing and towards more movement, which is observed both during late extinction and when processing future threats (Wallis et al. 2017, Alexander et al. 2020). Thus, it is critical to understand how IL interactions with subcortical and cortical structures that govern cognitive and autonomic function affecting extinction.

Importantly, prefrontal projections to the basolateral complex of the amygdala (BLA) were identified as a key input that undergoes synaptic plasticity during extinction learning (Amano et al. 2010, Cho et al. 2013), and is critical for extinction memory retrieval (Likhtik et al. 2008, Bukalo et al. 2015, Bloodgood et al. 2018, Hagihara et al. 2021). Accordingly, inhibiting IL projections to the BLA during extinction learning doesn’t impair extinction acquisition but hampers its retrieval, indicating that IL inputs to the BLA mediate extinction memory consolidation (Laurent and Westbrook 2009, Bukalo et al. 2015, Do-Monte et al. 2015, Bloodgood et al. 2018). However, non-specific pharmacological inhibition of the IL at the beginning of extinction also impairs within-session extinction acquisition (Sierra-Mercado et al. 2011). Thus, activity of IL pathways targeting regions other than the BLA may be important for modulating fear suppression during extinction learning.

The IL projects prominently to regions that control attention and autonomic activity, including cholinergic subnuclei of the basal forebrain (BF) such as the substantia innominata and the horizontal limb of the diagonal band of Broca (Room et al. 1985, Hurley et al. 1991, Zaborszky et al. 1997, Vertes 2004). Interestingly, BF subnuclei are implicated in modulating cued fear learning via projections to the amygdala and cortex (McDonald et al. 2011, Unal et al. 2015, Jiang et al. 2016, Aitta-Aho et al. 2018, Crimmins et al. 2023, Bratsch-Prince et al. 2024, Rajebhosale et al. 2024). Thus, we asked if IL input to the BF modulates fear extinction. First, we used retrograde tracing to compare prefrontal projections to the BF and BLA and showed that IL-BF output is overall denser than IL-BLA output. Then, using the immediate early gene cFos, we show that IL-BF Layer (L)5 projections are more active during extinction learning, whereas IL-BLA L2/3 projectors are more active during extinction retrieval. Further, using *in vitro* recordings, we demonstrate that L5 IL-BF projectors are most excitable at the end of extinction learning. Finally, we demonstrate that inhibiting the IL-BF pathway during extinction learning, impairs its acquisition but does not affect extinction memory retrieval the following day. Thus, we reveal a large output from deeper layers of the IL to the BF that aids in suppressing defensive fear expression during extinction learning but doesn’t partake in extinction memory formation. Collectively, our findings demonstrate, for the first time, that extinction is orchestrated by different IL pathways throughout learning and retrieval, whereby L5 IL-BF projections are upregulated during acquisition, whereas L2/3 IL-BLA projections are upregulated in retrieval.

## Materials and Methods

### Animals

Adult male C57BL/6J male mice (Jackson Laboratory), aged 9-11 weeks were group housed (2-4 per cage) under a 12 h light/dark cycle (lights on 8AM-8PM) with *ad libitum* access to food and water. All procedures were conducted under the regulation of the Hunter College Institutional Animal Care and Use Committee.

### Surgeries - Microinjections and Optic fiber implantations

For all surgeries, mice were anesthetized with 2% isoflurane in oxygen, placed in a stereotaxic frame (Kopf Instruments, Tujunga, CA) and maintained on 1.5% isoflurane throughout surgery (oxygen at a flow rate of 1L/min). Temperature was maintained at 37C±1C with a feedback-regulated heating pad. Mice received dexamethasone (1mg/mL, s.c.) and bupivacaine under the scalp (5mg/mL, s.c.) prior to incision.

For tracing/cFos experiments, two craniotomies were performed using a drill with a burr attachment. Mice were injected unilaterally in the right hemisphere with the 0.3µL of the retrograde tracers Cholera Toxin Subunit B (CTB), Alexa Fluor-488 and -647 (Invitrogen, Waltham, MA) into the BF (-0.15 mm AP,+1.5 mm ML,-4.5 mm DV from brain surface) and BLA (-1.6 mm AP,+3.15 mm ML,-4.1 mm DV from brain surface), counterbalanced by fluorophore and region, at a rate of 0.08μL/min with 10μL Hamilton syringes (QSW Stereotax Injector, Stoelting, IL). The craniotomies were then closed with bone wax, and the skin was closed with poly(glycolide-coacaprolactone) monofilament absorbable sutures (Covetrus). Postoperatively, all mice received carpofen for pain relief (1mg/mL, i.p.) and were group housed in cages warmed by a heating pad until recovery. Mice were allowed to recover from surgery for at least 1 week prior to handling.

For the optogenetic manipulation experiments, mice (n=9/grp) were injected bilaterally with 0.15µL of the inhibitory anterograde virus rAAV5-hSyn-eArch3.0-EYFP or its matched control rAAV5-hSyn-EYFP (10 × 10^12^ vg/ml; UNC Vector Core, NC) into the IL (+1.6 mm AP,± 0.4 mm ML,-2.0 mm DV from brain surface) at a rate of 0.08μL/min with 10μL Hamilton syringes (QSI Stereotax Injector, Stoelting, IL). Custom-order ferrules with attached optic fibers (exposed fiber length, 6mm, Newdoon, China) were then implanted over the BF (-0.15 mm AP, ± 1.5 mm ML, - 4.4 mm DV from brain surface) and cemented onto the skull using both opaque C&B Metabond (Parkell, USA) and an additional layer of different-colored dental cement (Teets, Lang Dental, USA) for animal identification. Mice were allowed to recover on a heating pad, and then the virus was left to express for 4-5 weeks prior to handling.

For the *in vitro* recordings, n=16 mice were injected bilaterally with 0.3µL of the retrograde virus rAAV-retro-hSyn-eYFP (1 ×10^12^ vg/ml; UNC Vector Core, Chapel Hill, NC) into the BF (-0.15 mm AP, ±1.5 mm ML, -4.4 mm DV from brain surface) and the skin was closed with poly(glycolide-coacaprolactone) monofilament absorbable sutures (Covetrus). Mice were allowed to recover for 2 weeks.

### Behavioral Experiments

*Context A*: Animals underwent fear conditioning in a plexiglass chamber with aluminum walls and a stainless-steel rod floor capable of delivering scrambled foot shock (Med Associates, VT). Overhead lamps maintained light levels at ∼40 Lux and the conditioning box was cleaned with ethanol between animals.

*Context B*: Animals underwent extinction in a custom-made gray wood box (45cm length x 13cm width x 20cm height), with a smooth paper floor that was changed between animals. Light levels were maintained at ∼70 Lux.

Auditory cues were delivered via an audio speaker (ENV-224AM, MedAssociates) located in the wall of the chamber (Context A) or above the enclosure (Context B) at approximately the same height as in Context A. Mice were presented with a 2kHz pure tone throughout the protocol, except for one group that also received 8kHz pure tones, all tones delivered at 100dB. Behavior was recorded using an infrared OptiTrack camera and Neuromotive software running in conjunction with Central software (Blackrock Neurotech, Salt Lake City, UT). Timestamped video data were analyzed offline.

#### Behavioral Protocols

*Handling and Habituation*: Mice were brought to the behavioral-adjacent room and allowed to acclimate 1 hour before the experiments started each day. Mice were first handled by the experimenter for 5min. The next day, mice were first handled for 5min and, at least one hour later, they underwent habituation to Context B for approximately 8min. At least one hour later, they were habituated to Context A, where they were exposed to five trials of the 30sec long conditioned stimulus (CS), a 2kHz tone (ITI, 60-120s). Each CS consisted of 50ms pips (amplitude modulated with 25ms linear increase followed by 25ms linear decrease), delivered once per second for 30s, as reported previously (Stujenske et al. 2022).

For the experiments assessing IL-BF projector activity during extinction, on Day 1, mice were allocated to one of three groups: a tone control group that received the same numbers of tone-alone trials as the other groups but did not undergo associative learning, an extinction learning group, and an extinction retrieval group:

- *Tone Control*: Day 1: 5 trials of a 2kHz tone CS in Context A, Day 2: 20 trials of a 2kHz tone CS in Context B, Day 3: 10 trials of a 2KHz CS tone in Context B.
- *Extinction Learning*: Day 1: 5 trials of a 2kHz tone CS co-terminating with a 1s US (0.7mA scrambled electric footshock) in Context A, Day 2: 20 trials of a novel 8kHz tone in Context B, Day 3: 5 trials of the 8kHz tone followed by 5 trials of the 2kHz CS tone in Context B.
- *Extinction Retrieval*: Day 1: 5 trials of a 2kHz tone CS co-terminating with a 1s US (0.7mA scrambled electric footshock) in Context A, Day 2: 20 trials of a 2kHz tone CS in Context B, Day 3: 10 trials of a 2kHz tone CS in Context B.

All mice were sacrificed and perfused 90min after the 6^th^ tone was delivered on the last day of the protocol (Day 3) to quantify expression of the immediate early gene cFos in IL.

For the experiments that were testing IL-BF pathway excitability *in vitro*: Following handling and habituation, mice were divided into one of four groups.

- *Tone Control*: Day 1: five trials of 2kHz tone-alone exposure, Day 2: two 2kHz tone-alone trials, perfusion 10min later.
- *Early Extinction*: Day 1: five trials of 2kHz CS paired with US (0.7mA scrambled electric foot shock), Day 2: two 2kHz tone-alone trials of extinction learning, perfusion 10min later.
- *Late Extinction:* Day 1: five trials of 2kHz CS paired with US (0.7mA scrambled electric foot shock), Day 2: twenty 2kHz tone-alone trials of extinction learning, perfusion 10min later.
- *Extinction Retrieval*: Day 1: five trials of 2kHz CS paired with US (0.7mA scrambled electric foot shock), Day 2: twenty 2kHz tone-alone trials of extinction learning, Day 3: two 2kHz tone-alone trials of extinction retrieval, perfusion 10min later

For experiments with optogenetic inhibition of IL inputs to the BF, handling and habituation were as described above, with the exception that all mice were also exposed to five trials of 35sec laser stimulation during Habituation to Context B. Then, mice underwent Fear Conditioning (context A), receiving five CS-US paired trials where the 2kHz CS delivery co-terminated with a 1s, 0.7mA footshock US. The next day, during Extinction Acquisition (Context B), mice received 10 trials of the 2kHz CS alone, coupled with a green laser (532nm, continuous stimulation, ∼10mW per hemisphere, Laserglow Technologies, Ontario), with the laser ramp-modulated during onset and offset. The next day, during Extinction Retrieval (Context B), animals were exposed to 10 trials of the 2kHz tone-alone.

For all behavioral analyses, time spent showing defensive freezing was manually quantified by an experimenter blind to group. The scoring consisted of measuring the amount of time spent freezing during the 30s prior to trial 1 (baseline), and during the 30s of all CS presentations.

### Tissue collection and immunostaining

For the neuroanatomical tracing and cFos immunostaining experiments, mice were deeply anesthetized with a mixture of ketamine (100mg/kg, i.p.) and xylazine (2 mg/kg, i.p.) and transcardially perfused with cold phosphate-buffered saline (PBS), followed by 4% paraformaldehyde (PFA) in PBS 90 minutes after the onset of CS #6 on Day 3. Brains were extracted and post-fixed in 4% PFA overnight. After cryoprotection in 30% sucrose in PBS, 40-micron histological sections were prepared on a cryostat (Cryostar) to evaluate: a) BF and BLA injection sites, b) cFos+ and CTB+ mPFC cell bodies. Mounted BF and BLA sections were coverslipped with ProLong Gold plus DAPI antifade mounting medium (ThermoFisher Scientific, MA) and imaged with a fluorescent episcope (Olympus BX53) to identify CTB injection placements. Tracing and cFos analyses were only carried out in mice with correct placements in BF or BLA and visible CTB+ cells in the mPFC (see **Fig. 1**). Immunohistochemistry was performed on mPFC sections at three defined points: Bregma AP +2.0mm, +1.8mm, +1.6mm.

**Figure 1:**
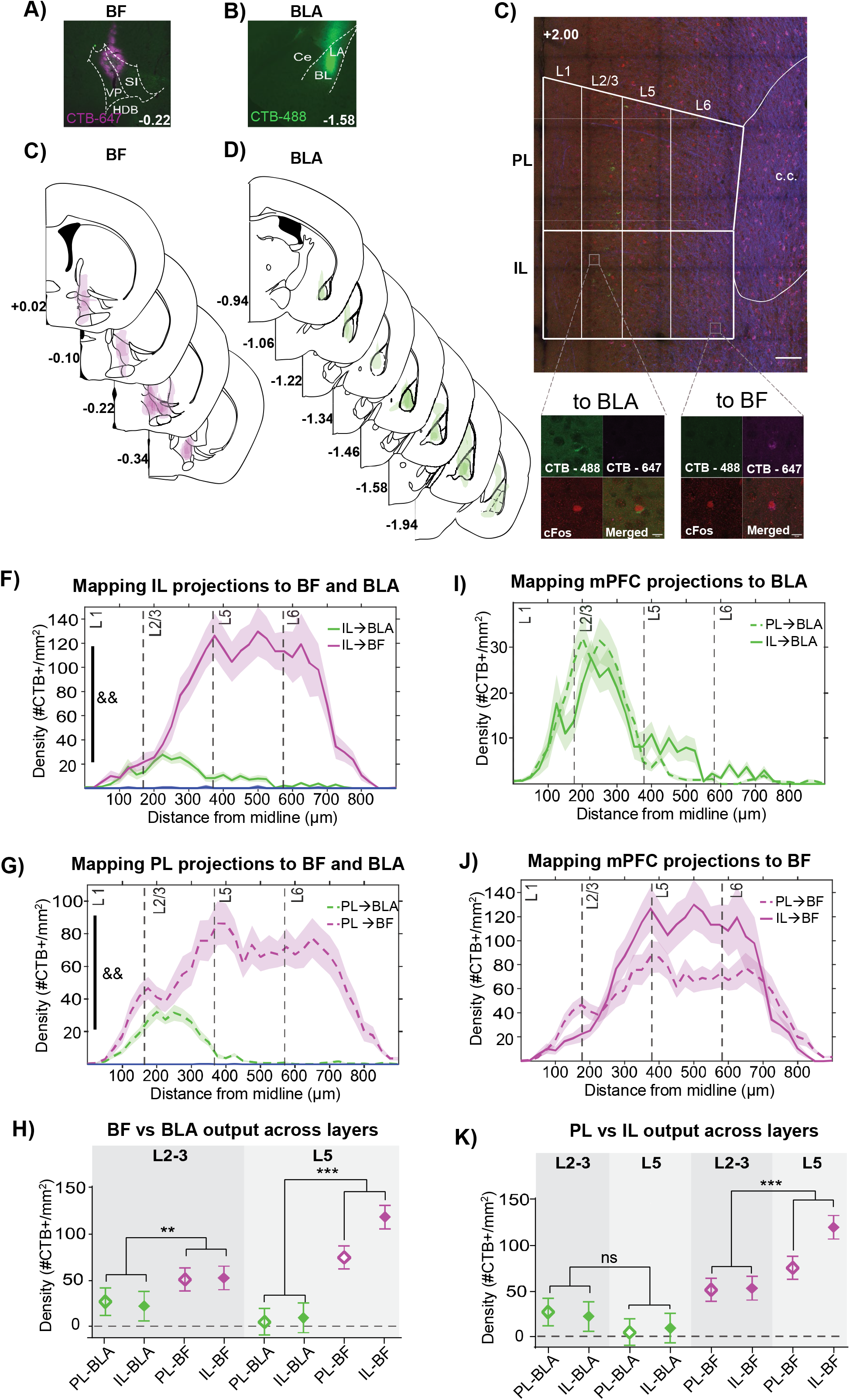
Comparative distribution of mPFC projections to the BF and BLA across cortical layers. **(A-B)** Examples of CTB injections in the BF and BLA. **(C-D)** Mapping of the full extent of CTB injections in the BF and BLA. **E)** Example of CTB and cFos labeling in the mPFC. Scale bar, 100μm. Insets: *Left,* example of CTB-488 labeling an IL-BLA L2/3 projector, which was also cFos+. *Right*, an example of CTB-647 labeling of an IL-BF L6 projector, which was also cFos+. Scale bar, 10μm. **F)** Comparative density mapping shows a significantly denser IL output to the BF than BLA (&&, post-hoc comparison of cortical region x subcortical target, p<0.01). **G)** Comparative density mapping shows a significantly larger PL output to the BF than BLA (&&, post-hoc comparison of cortical region x subcortical target, p<0.01). **H)** Mixed-model comparing density of BF and the BLA output shows denser PL and IL projections to the BF from both L2/3 and L5). **I)** Comparative density mapping shows that PL and IL projections to the BLA are similarly dense, with BLA projections from both peaking in L2/3 but spreading to deeper layers as well. **J)** Comparative density mapping and post-hoc comparisons show a significantly denser IL than PL output to the BF (&, p=0.028) arising from L5 (&&, p<0.01). **K)** Mixed-model comparing densities of PL and IL output across layers. Post-hoc comparisons show that there are no differences in PL vs IL outputs to the BLA across layers. However, there is a significantly denser projection from L5 IL than PL to the BF (&&, p<0.01). *Note that mPFC-BLA projections are illustrated with green (CTB-488) and mPFC-BF projections are illustrated with magenta (CTB-647) for visualization purposes only. During experiments, CTB-488 and CTB-647 injections were counterbalanced between BF and BLA.* Main effects: *, p<0.05; **, p<0.01; ***, p<0.001. Post-hoc comparisons: &, p<0.05, &&, p<0.01. All data shown as mean ±SEM. BF, basal forebrain; BL, basolateral nucleus; BLA, basolateral amygdala; Ce, central nucleus; HDB, horizontal limb of the diagonal band of Broca; IL, infralimbic cortex; LA, lateral nucleus; PL, prelimbic cortex; SI, substantia innominata; VP, ventral pallidum.

Free-floating sections were subsequently washed in 1x PBS (3x 5min), blocked in 5% normal donkey serum (NDS, Jackson ImmunoResearch) in PBSTriton 1% (Sigma Aldrich) at room temperature (1h), and then incubated overnight in Rabbit anti c-Fos antibody (1:2000, Abcam, #ab190289) and Mouse anti-myelin basic protein (MBP, 1:1000, Biolegend, #808401) in blocking solution at 4C, staining the corpus callosum to visualize the width of the mPFC. The next day, sections were washed in PBS (3x, 5min) and incubated in Donkey anti-Rabbit Alexa Fluor 594 secondary antibody (1:500, Life Technologies #A21207) and Donkey anti-Mouse Alexa-405 (Invitrogen, #a48257) in blocking solution for 2hr at room temperature. Sections were subsequently washed in 1xPBS (3x, 5min) before being mounted and coverslipped with ProLong Gold antifade mounting medium (ThermoFisher Scientific, MA).

### Cell counting

The three mPFC slices were imaged on a confocal microscope (40x, Leica SP8) with a Z-stack of six optical slices (3-4 μm/slice). Images were analyzed in ImageJ (NIH, Bethesda, MD), with each z-stack analyzed as a maximum intensity projection, with regions of interest (ROI) for IL and PL extracted for each slice, based on stereotaxic coordinates (Franklin and Paxinos 2013). For each ROI, fluorescing cells were counted using the Multipoint tool, and the accompanying x and y coordinates for each cell were saved. The coordinates were analyzed with custom written scripts (Matlab) that used the x,y coordinates of each cell to locate it in the mPFC. The x-axis was binned (25µm bins) and densities of cells per bin were calculated for each cell type. To calculate layer-specific parameters, bins encompassing 175-375μm from the surface were averaged into layers 2-3 (L2/3), and bins spanning 376-550μm from cortical surface were averaged as L5 for each animal (Little and Carter 2012).

### Slice Preparation

Slice preparation was performed as previously described (Friedman et al. 2016). Artificial cerebrospinal fluid (aCSF) was prepared in the following ion concentration; in (mM), NaCl 128; D-Glucose 10; NaH2PO4 1.25; NaHCO3 25; MgCl2 2; KCl 3; CaCl2 2. aCSF was ice-cold and oxygenated with 95% oxygen and 5% carbon dioxide. 10 minutes following the behavioral timepoint of interest, mice were anesthetized with isoflurane (1-chloro-2,2,2-trifluoroethyl-difluoromethylether). After confirming that the mouse was deeply anesthetized, an incision was made on the chest. Ice-cold oxygenated aCSF was trascardially perfused prior to rapid decapitation. After harvesting, the brain was blocked into mPFC-containing and BF-containing block. The mPFC-containing block was fixed on the buffer tray of a Microslicer (Microslicer DTK-1000, Dosaka EM, Kyoto, Japan) and the BF-containing block was soaked in 4% PFA in 1× PBS for placement evaluation. Acute brain slices containing mPFC neurons were cut at 250 μm-thick in cold oxygenated sucrose aCSF [in (mM), sucrose 227; D-Glucose 10; NaH2PO4 1.25; NaHCO3 24; MgCl2 2; KCl 3; CaCl2 2] using the microslicer. These slices were then transferred to a recovery chamber with oxygenated aCSF for 1 h at 36°C. The recovery chamber was then moved to room temperature with continuous oxygenation and slices were used for recording for up to a 4h period.

### Whole-cell patch-clamp recordings

Recordings were performed at 37°C using an inline solution heater (SH-27B) and temperature controller (TC-324C, Warner Instruments, MA). Slices were transferred to a recording chamber that was continually perfused with oxygenated aCSF at a flow rate of 3.0 ml/min. Recording pipettes were made from thick-walled borosilicate glass (BF150-86-10, Sutter Instrument, CA). Glass pipets were pulled by P-97 Flaming/Brown micropipette puller (Sutter Instrument, CA).

Patch pipet for whole-cell voltage-clamp and current-clamp (3–8 mΩ) was filled with internal solution [in (mM), K-gluconate 115; KCl 20; MgCl2 1.5; Phosphocreatine 10; K-ATP 2; Na-GTP 0.5; HEPES 10; pH 7.4, 284 mOsm]. IL was identified by anatomical location guided by anterior commissure spanning both hemispheres and lateral ventricles, and visualized with 4× objective lens (PLN 4X, Olympus, Japan). IL neurons were visualized under infrared light with 40× objective (LUMPLFLN, Olympus, Japan) immersed in aCSF. eYFP-labeled neurons were visualized with a fluorescent lamp (X-Cite 120Q, Lumen Dynamics, ON, Canada) with a 470nm filter and recordings were made from eYFP labeled neurons. Neurons of interest were identified by the presence of fluorescence in the soma. Neurons without fluorescence were not recorded. After the creation of a giga-Ω seal, the cell membrane was ruptured by small suction to create a whole-cell configuration. Different measurements of excitability were taken ∼2 minutes following the establishment of the whole-cell configuration in current-clamp mode (I=0): a) resting membrane potential (RMP), b) rheobase, i.e. as the minimum amount of current required to fire an action potential using a current ramp, 3) the relationship between increasing steps of current and the number of action potentials fired. For the latter, voltage responses to depolarizing current were recorded from 0pA to +200pA for 200 ms in increments of 10pA. To control for differences in RMP, current-injection protocols were performed at both RMP and -70 mV. Signals were digitized using a Multiclamp 700 B amplifier (Molecular Devices, San Jose, CA) and data was acquired using Axon Digidata 1550 B (Molecular Devices, San Jose, CA). The number of spikes during depolarizing current injection was counted by event detection (Clampfit). Rheobase was counted as the minimum current injection needed to elicit an action potential during the excitability protocol.

### Experimental Design and Statistical Analysis

All analyses were performed with Prism 10 software except for the mixed-effects model that was constructed in SPSS (version 25). For the anatomy analysis of mPFC projector distributions, we ran a mixed model with fixed effects of Projector (BF, BLA), mPFC Subregion (PL, IL), and Cortical Layer (L2/3, L5) and a random intercept for subject. For the cFos analysis, 2-way repeated measures ANOVAs were used to study differences across groups for the 2 layers (repeated measures) for each pathway, and 2-way ANOVAs to study differences between BF- and BLA-projectors across groups for L2/3 and L5. Due to variability in CTB tracer expression between cohorts, outlier analyses were run on the anatomical tracing experiment, using GrpahPad (Prism 10), which resulted in removal of two animals with abnormally high labeling through layers. For the behavioral analyses, a two-way repeated measures ANOVA (two-way rmANOVA) was performed with “Group” and “Trial” as factors and Tukey’s or Sidák’s as the posthoc test. For comparisons between specific behavioral trial-bins, RMP, and rheboase across groups, unpaired t-tests were performed in normal distributions or Mann-Whitney tests if the distributions failed to pass the normality test. Differences in number of action potentials (APs) between groups were investigated using a two-way repeated measures ANOVA with “Current Step” and “Group” as factors. All data are expressed as mean ± SEM and significance was defined as p<0.05.

## Results

The mPFC has a strong reciprocal connection from L2/3 with the BLA, which partakes in fear and extinction processing (Quirk et al. 2003, Little and Carter 2012, Arruda-Carvalho and Clem 2014, Burgos-Robles et al. 2017, Klavir et al. 2017, Bloodgood et al. 2018). Intrigued by these connections, we aimed to compare the distribution of IL projections to the BF with those to the BLA, and to determine whether IL projections to the BF and BLA are similarly upregulated with extinction. Previous work in rats, using the anterograde tracer *Phaseolucoagglutinin* (Room et al. 1985, Hurley et al. 1991, Vertes 2004), showed that the IL has a stronger projection to the BF than the PL. Thus, we were also interested in understanding whether this pattern is retained in mice. To answer these questions, we injected the retrograde tracer CTB in the BF and BLA and assigned mice to one of three behavioral conditions, Tone-Control, Extinction Learning, or Extinction Retrieval (behavior in figures 2 and 3). All groups of animals were transcardially perfused at the specified time points. We then mapped the distribution of CTB-labeled mPFC soma projecting to the BF and BLA (**Fig. 1**) and analyzed the overlap of CTB and cFos expression to investigate pathway-specific activity related to each behavioral condition (**Fig. 2-3**).

**Figure 2:**
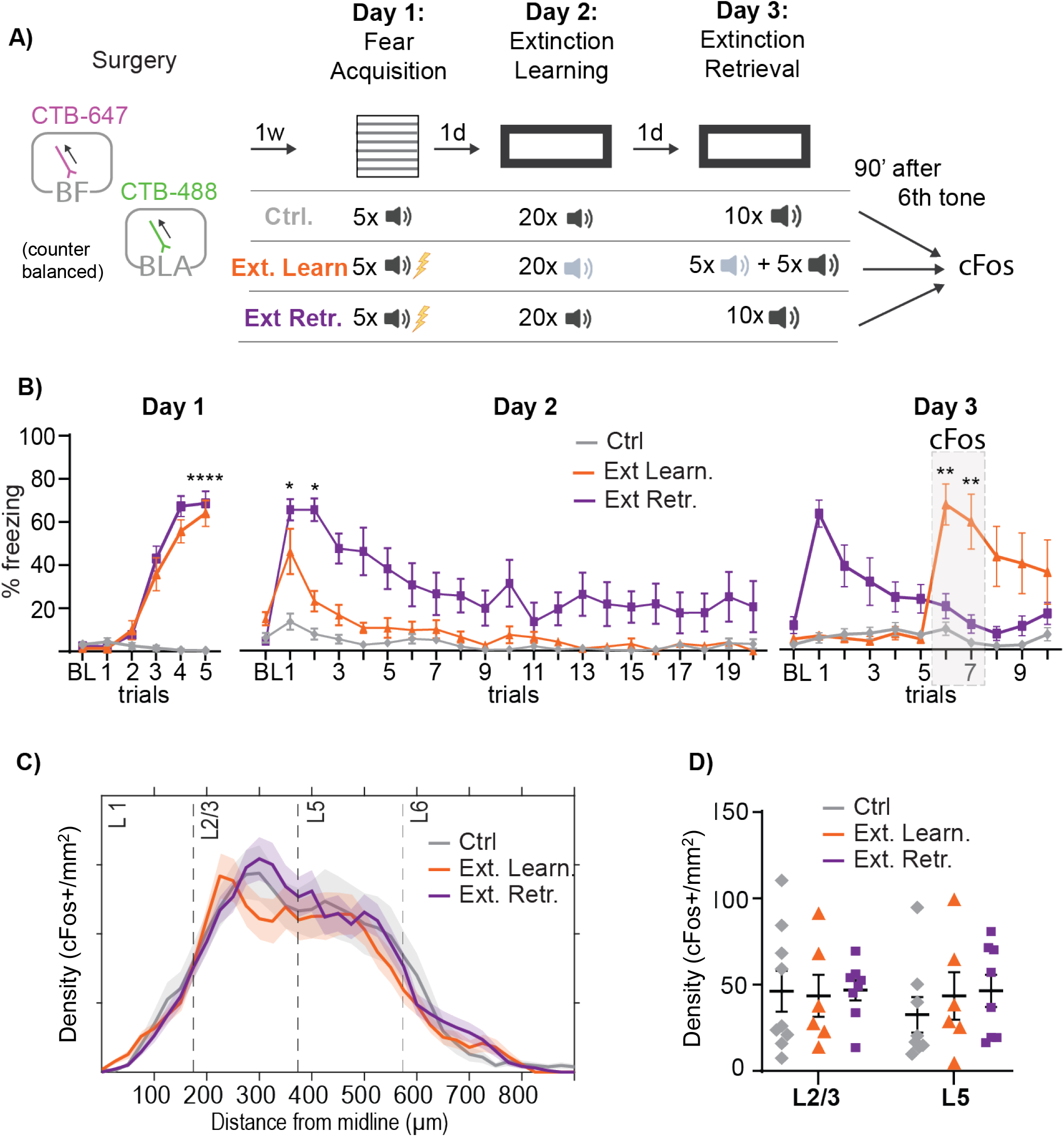
Global IL L2/3 and L5 activity is similar during extinction learning and retrieval. **A)** Timeline of CTB injection surgeries and behavioral paradigm tailored for cFos expression. One week after CTB injections in the BF and BLA, animals were randomly assigned to one of three groups. The Tone Control group (Ctrl, grey) was exposed to unpaired tones across Days 1-3. The Extinction learning group (Ext Learn., orange) was fear conditioned with five CS-US pairings on Day 1. Then, to control for tone exposure, this group received 20 trials of a new, unpaired tone on Day 2, and then on Day 3, another five trials of the unpaired tone, followed by five trials of the CS for extinction learning. The Extinction Retrieval group (Ext Ret., purple) was fear conditioned with five CS-US pairings on Day 1, extinguished with 20 CS trials on Day 2 and, on Day 3, underwent extinction retrieval with 10 CS trials. All animals were perfused 90-minutes after the 6th tone on Day 3. **B)** Percent defensive freezing in all groups throughout the behavioral paradigm. Day 3, grey box highlights the trials for timing cFos capture, when the Extinction Learning group freezing was significantly higher than both in Controls and Extinction Retrieval groups. **C)** Non-projection specific density mapping of cFos+ cells across all layers of the IL in all behavioral groups. **D)** There were no differences in the average number of IL cFos+ cells across groups in either L2/3 or L5. *, p<0.05; **, p<0.01; ***, p<0.001. All data shown as mean ±SEM.

**Figure 3:**
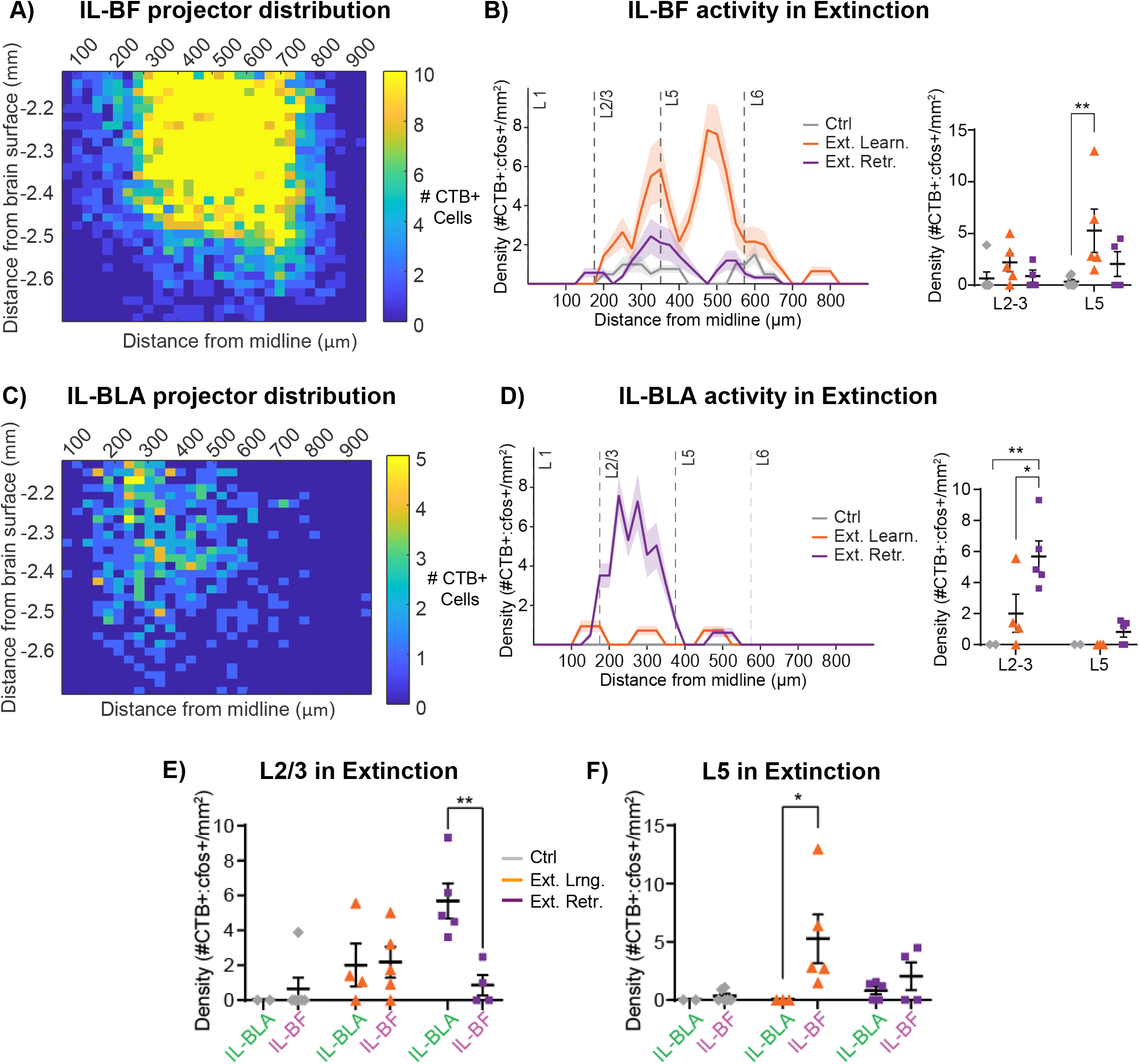
L5 IL-BF projectors upregulate during extinction learning, whereas L2/3 IL-BLA projectors upregulate during extinction retrieval. **A)** Heatmap showing the anatomical distribution of IL projectors to the BF along the medio-lateral and dorso-ventral axes of the IL. Color bar, average number of CTB+ cells. **B)** Cortical layer density map (left) and quantification (right) comparing distribution of cFos+ IL-BF projectors in the three behavioral groups across superficial and deep layers of IL. There are significantly more cFos+ IL-BF projectors in L5 in the Extinction Learning group than in Controls. **C)** Heatmap showing the anatomical distribution of IL projectors to the BLA along the medio-lateral and dorso-ventral axes of the IL. Color bar, average number of CTB+ cells. **D)** Density map (left) and quantification (right) comparing distribution of cFos+ IL-BLA projectors in the three behavioral groups across superficial and deep layers of IL. There are significantly more cFos+ L2/3IL-BLA projectors in in the Extinction Retrieval group than in Controls and in Extinction Learning groups. **E)** Comparison of L2/3 IL-BF vs IL-BLA pathway activity in behavior. The L2/3 IL-BLA pathway is significantly more active than the IL-BF pathway in the Extinction Retrieval group. F) Comparison of L5 IL-BF vs IL-BLA pathway activity in behavior. The L5 IL-BF pathway is significantly more active than the IL-BLA pathway in the Extinction Learning group. *, p<0.05; **, p<0.01; ***, p<0.001.

### IL and PL output from superficial and deep layers is denser to the BF than to the BLA

To compare the distribution of mPFC projectors in BF and BLA pathways, we injected C57B/6J male mice with the retrograde tracer CTB (0.3μl) in the posterior substantia innominata/ ventral pallidum region of the BF (-0.15 mm AP, +1.5 mm ML, -4.5 mm DV from brain surface) and in the BLA (-1.6 mm AP, +3.15 mm ML, -4.1 mm DV from brain surface, **Fig. 1A-B**). The CTB was counterbalanced for fluorophores (CTB-488 or CTB-647) at each injection site. All sites were checked for correct targeting, and those with incorrect placements were removed from analysis, with the resulting n=16 BF and n=11 BLA (**Fig. 1C-D**). The density and distribution of CTB-expressing soma in the mPFC were analyzed and averaged across three anterior-posterior locations (Bregma +2.0mm, +1.8mm, +1.6mm). At each location, the x-coordinates of CTB-expressing soma were binned (25 μm bins) and mapped along the medio-lateral axis of the mPFC, spanning from the pial surface to the corpus callosum, and then averaged across superficial (L2/3, 175-374μm from midline) and deep layers (L5, 375-550μm from midline, **Fig. 1E**).

Using a mixed model analysis to account for mPFC cortical region (PL vs IL), subcortical projection target (BF vs BLA) and Layer (L2/3 vs L5), we first identified a significant effect of subcortical projection target (F_(1,129)_=54.36, p<0.001), with a denser mPFC projection to the BF than to the BLA (BF: 75.1 cells/mm^2^ ±8.8 vs BLA: 7.6 cells/mm^2^ ±10, p<0.001), with both the IL and the PL showing the same pattern (cortical regions x subcortical target, F_(1,110)_=0.64, p=0.43, **Fig. 1F-H**). Next, we looked at the density of subcortical innervation from different cortical layers, and saw a significant difference in density of BF vs BLA innervation from superficial vs deep layers (subcortical target x layer, F_(1,109)_=14.79, p<0.001). A pairwise comparison showed that mPFC (IL and PL) projections to the BF are denser than those to the BLA out of layers L2/3 (p=0.004) and out of L5 (p<0.001, **Fig. 1H**). Further, the BF receives denser input from mPFC L5 (p<0.001) but there was enough variability in mPFC output to the BLA, such that there were no differences in mPFC-BLA output between L2/3 and L5 (p=0.17, **Fig. 1K**).

### Overall output is denser from IL than PL

Next, we were interested in whether mice show a denser projection from the IL than the PL to the BF, as was previously observed in rats (Vertes 2004). The mixed model analysis showed a threshold effect of cortical region (PL vs IL, F_(1,110)_=3.92, p=0.05), with an overall denser projection from the IL than the PL across subcortical targets (IL: 49.5 cells/mm^2^ ±9, PL: 33.2 cells/mm^2^ ±9). Despite the overall denser projection to the BF than the BLA (subcortical projection target, F_(1,129)_=54.36, p<0.001), there was no significant cortical region x projection target interaction (F(_1,110_)=0.64, p>0.05, **Fig. 1I-K**), indicating that projections from the IL are denser than those of the PL overall. Note that although L5 IL-BF projections were seemingly more numerous than L5 PL-BF projections (**Fig. 1J-K**), the three-way comparison of cortical region x subcortical target x layer was not significant (F_(1,109)_=1.007, p>0.05).

Interestingly, the number of BF and BLA co-projectors was very sparse for both IL and PL, suggesting that mPFC projections to the BLA and the BF are largely non-collateralizing. Thus, overall, our anatomical analyses show that 1) the mPFC sends a denser projection to the BF than to the BLA from both superficial and deep layers, 2) the mPFC-to-BF projection is denser from deeper than superficial layers, 3) projections from the IL are denser than from the PL to both subcortical targets, 4) mPFC projections to the BLA and BF have minimal collateralization.

### L5 IL-BF projectors are more active during extinction learning whereas L2/3 IL-BLA projectors are more active during extinction retrieval

Having established a differential distribution of IL pathways to the BF and the BLA, we were interested in identifying whether these pathways have a similar activity profile during extinction. To do so, the animals that were injected with CTB were divided into three groups Tone Control (n=9), Extinction Learning (n=9), and Extinction Retrieval (n=10), based on their behavioral condition during cFos capture (**Fig. 2A**). Mice first underwent handling and habituation, and then were exposed to a 3-day paradigm, where on Day 1 they were either exposed to five 2kHz tone-alone trials (Tone Control), or fear conditioned with five paired 2kHz CS-US trials (Extinction Learning and Retrieval groups). On Day 2, mice were either exposed to 20 2kHz tone-alone control trials (Tone Control group), 20 trials of a novel neutral tone (8kHz) to control for total trial exposure (Extinction Learning group), or 20 2kHz tone-alone trials for fear extinction (Extinction Retrieval group). Then, on Day 3, animals in the Tone Control group were exposed to 10 2kHz tone-alone trials. Mice in the Extinction Learning group were first exposed to five trials of the neutral 8kHz tone to control for total trial exposure, followed by 5 trials of the fear conditioned 2kHz CS to capture extinction learning. Mice in the Extinction Retrieval group were exposed to 10 trials of the previously extinguished 2kHz CS to capture extinction retrieval. Animals were perfused 90-min after trial 6, targeting cFos expression to the start of acquisition in the Extinction Learning group, to low freezing due to good retrieval in the Extinction Retrieval group and to low freezing due to absence of fear learning in the Tone Controls (**Fig. 2A**).

Fear conditioning (**Fig. 2B**) resulted in significant differences in freezing across groups (F(2,125)=57.99, p<0.0001) and a trial by group interaction (F(10,125)=29.09, p<0.0001), with the two fear conditioned groups freezing more than Tone Control group by the end of the session (trial 5, Control 0%±00 vs Extinction Learning group 64.1%±6, Extinction Retrieval group 68.7%±5.6, both p<0.001). Importantly, the two fear conditioned groups showed similar levels of fear by the end of training (trial 5, Extinction learning 64.1%±6 vs Retrieval group 68.7%±5.7, p=0.84). Then, on Day 2 (**Fig. 2B**), there were significant differences between groups (F(2,25)=7.25, p=0.003) with a trial x group interaction (F(40,500)=3.88, p<0.0001).

Post-hoc comparisons showed that the controls did not freeze to the tone, whereas mice that underwent 20 extinction trials significantly decreased freezing during the session (trial 1 vs. 20, p= 0.02). Notably, animals that were exposed to 20 trials of a new tone on Day 2, showed some freezing at the beginning of the session, which was likely due to novelty or fear generalization (trials 1-2 Controls vs Extinction Learning, p=0.04), but that this response was gone by trial 3, when their freezing was the same as controls (p=0.13), staying low for the rest of the session.

On Day 3, the Tone Control group continued to show low freezing throughout the session, whereas the Extinction Learning group also showed low freezing during the neutral tone presentations (trials 1-5), but then increased freezing when exposed to the CS starting with trial 6 (**Fig. 2B**), when they froze significantly more than both the Control (Extinction Learning 67.1%±9.7 vs Tone Controls 9.5%±3.1, p=0.006) and the Extinction Retrieval group (Extinction Learning 67.1%±9.7 vs Retrieval 21.2%±5, p=0.014), whereas the Control and Extinction Retrieval groups showed similarly low freezing (p=0.29). The Extinction Learning group then decreased freezing such that it was no longer different from the Extinction Retrieval or Control groups by trial 10 (p>0.05 for both). Thus, our cFos timing (trial 6), aimed to capture the Extinction Learning group in a relatively higher fear state than both the Control and Extinction Retrieval groups, which were in a similarly low fear state by comparison.

Turning to neural activity, we first asked whether overall, the IL is differentially active during extinction learning or retrieval versus the control condition. To address this question, we compared the density of cFos+ cells between superficial and deep layers of the IL and found no differences in overall density of active cells in the IL across behavioral groups (F(_2,20_)=0.0178, p=0.982) or layers (F(_1,20_)=0.0004, p=0.984, **Fig. 2C-D**). This finding shows that when considered overall, IL activity doesn’t change across these behavioral states.

Next, we took advantage of retrograde tracers to identify pathway-specific activity in the IL during extinction. IL-BF pathway tracing showed a large projection peaking in L5 (**Fig. 3A**, see also **Fig. 1F,J**), and thus we were interested in whether there was evidence of it being active in these layers during extinction. In the IL-BF pathway, a group by layer repeated measures ANOVA showed a significant effect of group (F(_2,12_)=4.98, p=0.02). Tukey’s multiple comparisons revealed that the density of active IL-BF projectors was significantly higher in the Extinction Learning than the Tone control group (p<0.01), whereas the Extinction Retrieval group did not differ in activity from Tone controls (**Fig. 3B**, p=0.13). This increase in IL-BF activity was only observed in L5, without any significant differences in L2/3 IL-BF activity across behavioral groups (p>0.05, **Fig. 3B**).

Turning to the IL-BLA pathway, the overall anatomy showed a relatively less dense projection, peaking in L2/3 but also distributing to deeper layers (**Fig. 3C**, see also **Fig. 1F,I**). In regards to behavior-dependent activity in the IL-BLA pathway, we found significant main effects for both layer (F(1,8)=9.73, p=0.01) and group (F(2,8)=6.77, p=0.02), as well as a layer by group interaction (F(2,8)=4.38, p=0.05). Tukey’s post-hoc tests revealed a significantly higher density of active L2/3 IL-BLA projectors in the Extinction Retrieval group compared to Tone Controls (p<0.01) and compared to the Extinction Learning group (p<0.01, **Fig. 3D)**. However, L5 IL-BLA projectors did not differ across groups (p>0.05). Thus, IL-BLA L2/3 projections are more active during extinction memory retrieval.

Finally, we were interested in how activity in these two IL pathways compared across layers during behavior. In IL L2/3 efferents, a two-way ANOVA showed an interaction between IL pathway x group (F(2,21)=3.774, p=0.04). A Sidák posthoc test showed that during Extinction retrieval, the L2/3 IL-BLA pathway was significantly more active than the IL-BF pathway (p=0.03, **Fig. 3E**). An analysis of L5 showed a main effect of pathway (F (1, 19) = 4.94, p=0.04), with Sidák posthoc test revealing significantly higher L5 IL-BF than IL-BLA projector activity during Extinction Learning (p=0.02, **Fig. 3F**). In sum, this experiment shows that the L5 IL-BF pathway becomes more active during extinction learning, whereas the L2/3 IL-BLA pathway is upregulated during extinction retrieval.

### The IL-BF pathway gains excitability with extinction learning and becomes less excitable at extinction retrieval

The finding that the L2/3 IL-BLA pathway is more active during extinction retrieval compared to extinction learning (**Fig. 3B**) is in line with previous work showing that inhibition of this pathway during extinction learning doesn’t affect learning but impairs retrieval (Bukalo et al. 2015, Bloodgood et al. 2018). Further, *in vitro* IL recordings show that it becomes more excitable in the hours after extinction (Santini et al. 2008, Cruz et al. 2014), with cells likely undergoing cellular and molecular processes of consolidation that allow for the IL to act via target structures such as the BLA (Santini et al. 2004, Do-Monte et al. 2015, Bloodgood et al. 2018).

Given our cFos finding that L5 IL-BF projectors were active during fear extinction learning, we were interested in whether excitability changes with extinction in the IL-BF pathway. Despite timing perfusions for cFos assessment to the beginning of extinction learning (**Fig. 2-3**), cFos transcription has a relatively low temporal resolution, making it difficult to know whether IL-BF is more likely to be excited early or later in extinction acquisition. To address these questions, mice were injected with the retrograde virus AAV2-hSyn-eYFP (UNC Vector Core) in the BF and then *in vitro* patch recordings were obtained at resting membrane potential (RMP) and when the membrane was held at -70mV from identified IL-BF projectors in mice that were either exposed to tone alone, early or late extinction learning, or extinction retrieval (**Fig. 4A-B**). To do so, after two weeks of viral expression, mice were handled and habituated to the context, followed by five trials of tone-alone exposure (Tone Controls), or five trials of paired CS-US fear conditioning. The next day, *in vitro* recordings were obtained from IL-BF projecting cells in deeper IL layers, either after two trials of 2kHz tone-alone exposure (Tone Controls, n=14 cells from n=3 mice), two 2kHz tone-alone extinction trials (Early Extinction n=26 cells from n=5 mice), or twenty 2kHz tone-alone extinction trials (Late Extinction, n=16 cells from n=3 mice). A fourth group of animals went through twenty 2kHz tone-alone trials of extinction learning, and then the next day was exposed to two 2kHz tone-alone extinction retrieval trials prior to *in vitro* patch recordings (Extinction Retrieval, n=28 cells from n=5 mice).

**Figure 4:**
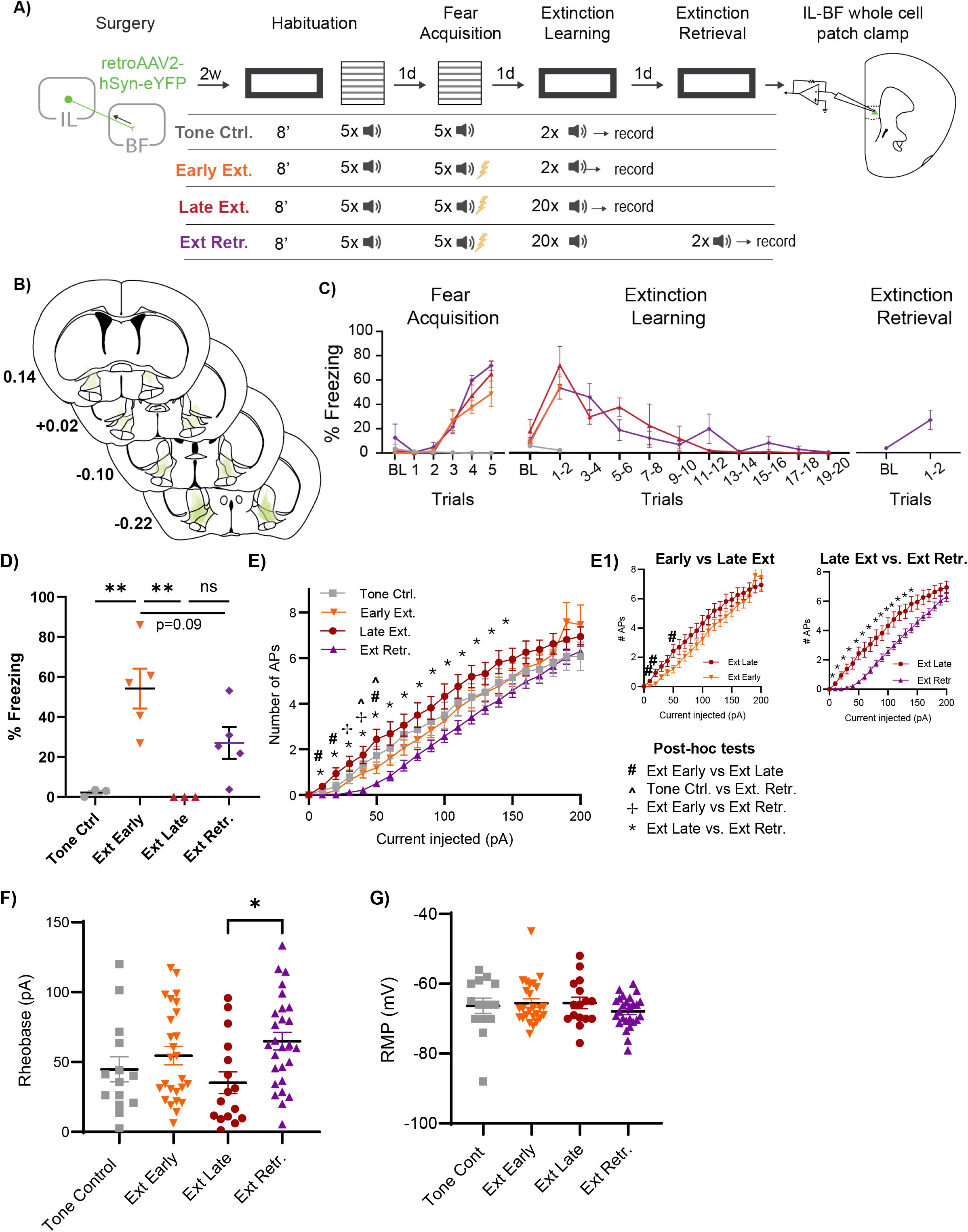
Deep layer IL-BF projectors become more excitable at the end of extinction learning. **A)** Timeline of injection surgeries and behavioral paradigm prior to *in-vitro* recordings of IL projectors to the BF. IL-BF projectors were identified with injections of rgAAV2-hSyn-eYFP. After 2 weeks of expression, animals were randomly assigned to one of four groups to probe IL-BF projector excitability during Early Extinction (Early Ext., orange), Late Extinction (Late Ext., red), Extinction Retrieval (Ext. Retr., purple), or Tone Control (Tone Ctrl., grey). Mice were sacrificed for *in vitro* recordings 20-min after the last behavioral trial. **B)** Anatomical mapping of viral injections in the BF. **C)** Percentage defensive freezing for all behavioral groups. Vertical arrows mark the last two trials of behavior for each group, after which the mPFC was sliced for *in vitro* recordings. **D)** Average percent freezing in the last two trials before perfusion. **E)** Excitability curves at RMP shown by the number of action potentials (APs) in response to increasing injections of current for each behavioral group. Significant post-hoc tests are marked with their respective symbols shown in the key. **E1**) Insets comparing different groups re-plotted for clarity. All significance testing was done on four groups. *Left*, Early vs Late Extinction excitability curves, post-hoc tests that reached significance or trend-level significance are marked with #, indicating increased excitability in Late vs Early extinction in IL-BF projectors. *Right*, IL-BF projectors in Late Extinction are significantly more excitable than in Extinction Retrieval in a wide range of testing conditions. **F)** Rheobase (pA), or the lowest level of current to evoke an action potential, across groups. IL-BF projectors show significantly lower rheobase in Late Extinction than Extinction Retrieval. **G)** IL-BF resting membrane potential (mV) shows no difference across groups.

Freezing behavior on conditioning day was analyzed with a repeated measures ANOVA that tested freezing over trials in the four groups. This analysis showed a main effect of trial (F(_3.5,42.2_)=46.06, p<0.0001), group (F(_3,12_)=11.19, p<0.001), and a trial by group interaction (F(_15, 60_)=5.76, p<0.001). Subsequent multiple comparisons showed that by trial 5 of fear conditioning, the fear conditioned groups had higher freezing than Tone Control group (p<0.05 for each conditioned group, **Fig. 4C**), whereas the fear conditioned groups had similar amounts of defensive freezing (p>0.05), indicating that they had learned the CS-US association.

On extinction learning day, there was a significant effect of group (one-way ANOVA, (F(_3,12_)=4.78, p=0.02), and trial (F(_3.2,24.2_)=15.2, p<0.0001), indicating that animals learned to extinguish fear. A one-way ANOVA evaluating defensive freezing in all groups during the two trials preceding *in vitro* recordings showed a significant effect of group (p=0.002), with post-hoc comparisons showing that the Early Extinction group froze significantly more during the first two trials of extinction learning than Tone Control and Late Extinction groups (p<0.01), **Fig. 4C-D**). Patch recordings of IL-BF projector excitability at RMP showed a significant effect of group (F(_3,80_)=3, p=0.03), current step (F(_3,256_)=351, p<0.0001), and a current step x group interaction (F(_60,1600_)=1.96, p<0.0001). Interestingly, post-hoc comparisons showed that despite mice being in high vs low fear states for the Early Extinction vs Tone Control groups (**Fig. 4D**), respectively, the excitability of IL-BF cells at RMP in these groups was not different (**Fig. 4E**, Tone Controls: n=14 cells from N=3 mice, grey, Early Extinction: n=26 cells from N=5 mice, orange, p>0.05), indicating that early in extinction learning, the IL-BF pathway did not differ in excitability from the tone-alone control condition. Later in extinction learning, freezing for the Late Extinction group decreased, such that by trial 19-20 defensive freezing was significantly lower than during trials 1-2 (p<0.001), and was lower than trials 1-2 for the Early Extinction group (p<0.01, **Fig. 4C-D**). Interestingly, Tukey’s multiple comparisons revealed that in the Late Extinction group, IL-BF excitability at RMP increased relative to the Early Extinction group at several current levels (**Fig. 4E-E1**, 10mA p=0.08, 30mA p=0.05, 50mA, p=0.08).

The next day, the Extinction Retrieval group showed a trend towards significantly decreased freezing from Early Extinction (p=0.09), and no difference in freezing from Late Extinction group (p>0.5, **Fig. 4D**). However, during Extinction Retrieval, IL-BF excitability at RMP was significantly lower than in the Late Extinction group (**Fig. 4E-E1,** 10-140mA pulses, all p<0.05). IL-BF excitability during Extinction Retrieval was also lower than during Early Extinction and from Tone Control at several current-level steps in the lower, more physiological stimulation range (**Fig. 4E**, 10-50mA pulses). We then tested IL-BF projector rheobase, or the current needed to drive a cell to spike at RMP, in all behavioral groups. This analysis showed that rheobase was significantly different between behavioral groups (Kruskall-Wallis, p<0.05), with multiple comparisons revealing that less current was needed to drive an IL-BF cell to fire in the Late Extinction group than in the Extinction Retrieval group (p<0.05, **Fig. 4F**). These findings confirm that the IL-BF pathway is more excitable as extinction learning progresses, and then becomes less excitable during extinction retrieval. Notably, changes in excitability were not accompanied by any change in resting membrane potential (**Fig. 4G**, one-way ANOVA, F(_3,78_)=0.81, p=0.49) and there were no changes in excitability observed when IL-BF projectors were held at -70mV (data not shown), indicating that IL-BF projectors are likely to be more synaptically driven by the network during extinction learning rather than retrieval, without intrinsically changing these cells.

### IL-BF input promotes within-session extinction

Given that L5 IL-BF neurons upregulate their activity and become more excitable during extinction learning, we next wanted to know if this projection functionally contributes to extinction. To this end, we injected the inhibitory opsin AAV5-hsyn-eArch3.0-eYFP (n=9) or its control AAV5-hsyn-eYFP (n=9) in the IL and bilaterally implanted optic fibers over the BF (**Fig. 5A-B**). After 4-5 weeks of viral expression, animals were habituated to tone and 35s exposure to 532nm laser (3sec on and 2sec off ramps, (Mahn et al. 2016)), and then underwent fear conditioning with five CS-US pairings 24h later. The next day, animals underwent extinction learning with IL terminals to the BF inhibited during each CS, and extinction retrieval was tested the next day in the absence of laser (**Fig. 5C**). During habituation, mice in both groups similarly did not freeze to presentations of tone (eYFP 2.8%, eArch 2.2%, p>0.05) or laser (eYFP, 0.5%, eArch, 1.1%, p>0.05, data not shown). Then, during fear conditioning, both groups acquired the CS-US association similarly (**Fig. 5D**, repeated-measures ANOVA, trial (F(_2.72, 43.05_)=69.46, p<0.0001, group F (_1,16_)=0.64, p=0.44). Then, during extinction learning, laser was administered to both groups (eYFP and eArch) during the CS, which resulted in a significant difference in freezing between groups (**Fig. 5D**, repeated measures ANOVA, F(_1,16_)=6.45, P=0.02) as well as a group by trial interaction (**Fig. 5E**, F(_5,80_)=3.99, p=0.003). Post-hoc comparisons revelated that the eArch group was not different from the eYFP group at baseline or at the beginning of extinction (trial 1-2) but was freezing higher than the eYFP group by trials 3-4 of extinction learning (p<0.05), and was trending towards significant differences at trials 5-6 (p=0.09), indicating an impairment in within-session extinction (**Fig. 5E**). Notably, when tested on extinction retrieval the next day, there was no difference in freezing between groups (**Fig. 5D-E**, F (1,16) = 0.04, p>0.84)), indicating that the IL-BF pathway is important for the online expression of extinction learning, but not for storing the extinction memory.

**Figure 5:**
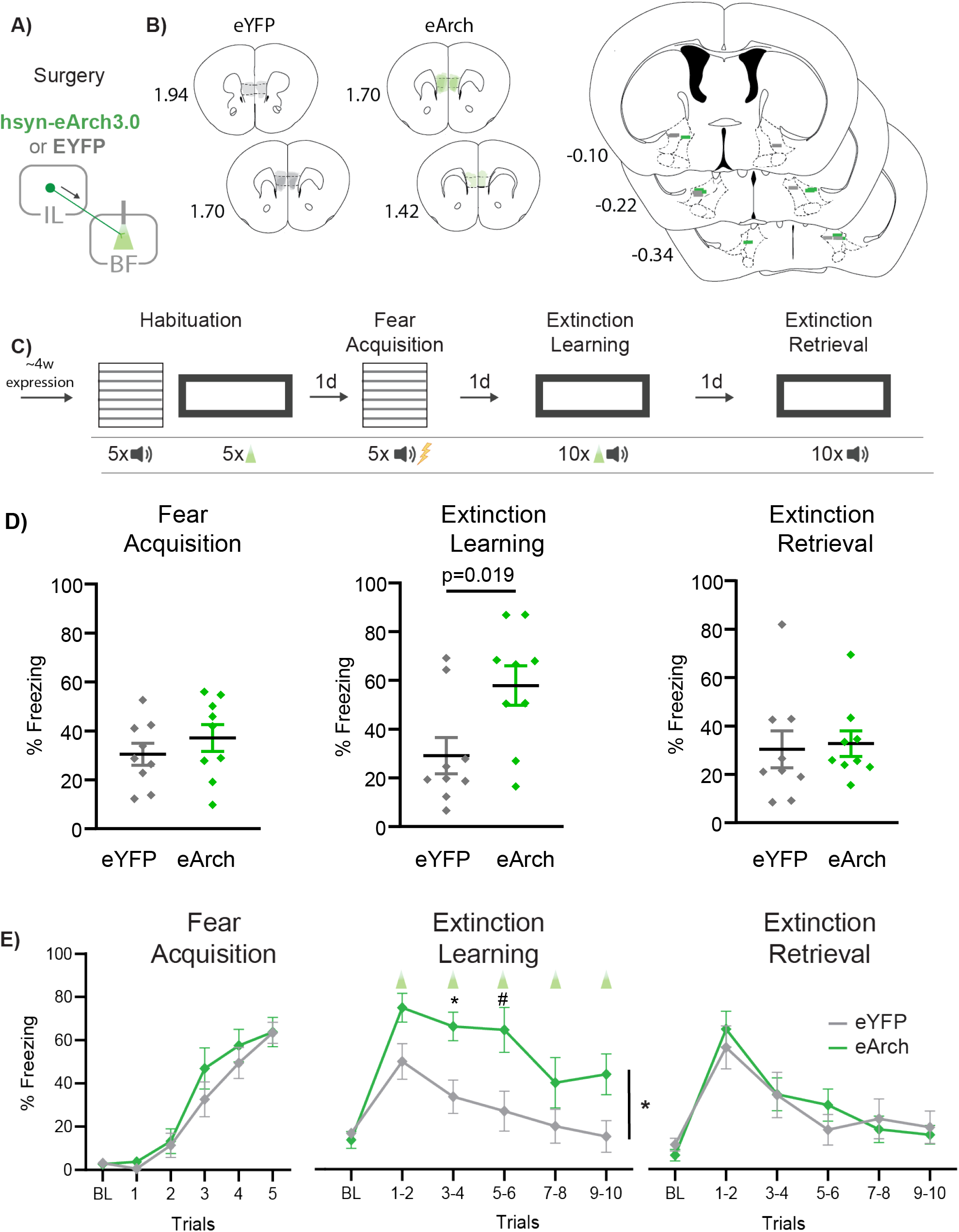
Inhibiting IL projections to the BF impairs extinction learning. **A)** Schematic showing the injections and fiber placement surgeries. **B)** Mapping of the viral spread of the virus in the IL (Left), and fiber placements in the BF (Right). Grey, eYFP; green, eArch. **C)** Schematic of behavioral paradigm. Mice were first habituated to the fear conditioning context and the tone, as well as to the extinction context and the laser. The next day, mice were fear conditioned with five CS-US pairings. The next day, mice underwent fear extinction for ten trials with laser light delivery during the tone, inhibiting IL inputs to the BF. One day later, mice were re-exposed to the extinction context during a 10-trial session of extinction retrieval. **D)** Average percent defensive freezing in eYFP and eArch groups across the entire behavioral session**. E)** Average percent defensive freezing across all trials, show in two-trial bins. Both groups learned the CS-US association similarly. However, inhibition of IL input to the BF significantly slowed extinction learning. The next day, both groups showed similar levels of freezing during extinction retrieval.

## Discussion

The IL is a critical region for extinction and safety learning (Milad et al. 2005, Giustino and Maren 2015, Kim et al. 2016, Felix-Ortiz et al. 2024, Ng et al. 2024), prompting us to investigate whether one of its major outputs, the pathway to the BF (Room et al. 1985, Hurley et al. 1991, Zaborszky et al. 1997, Vertes 2004), plays a role in extinction. Here, we show that the IL-BF pathway originates from L2/3 and L5, peaking at L5, that during extinction learning the L5 output becomes more active, and becoming as extinction learning progresses (**Fig. 6**).

**Figure 6:**
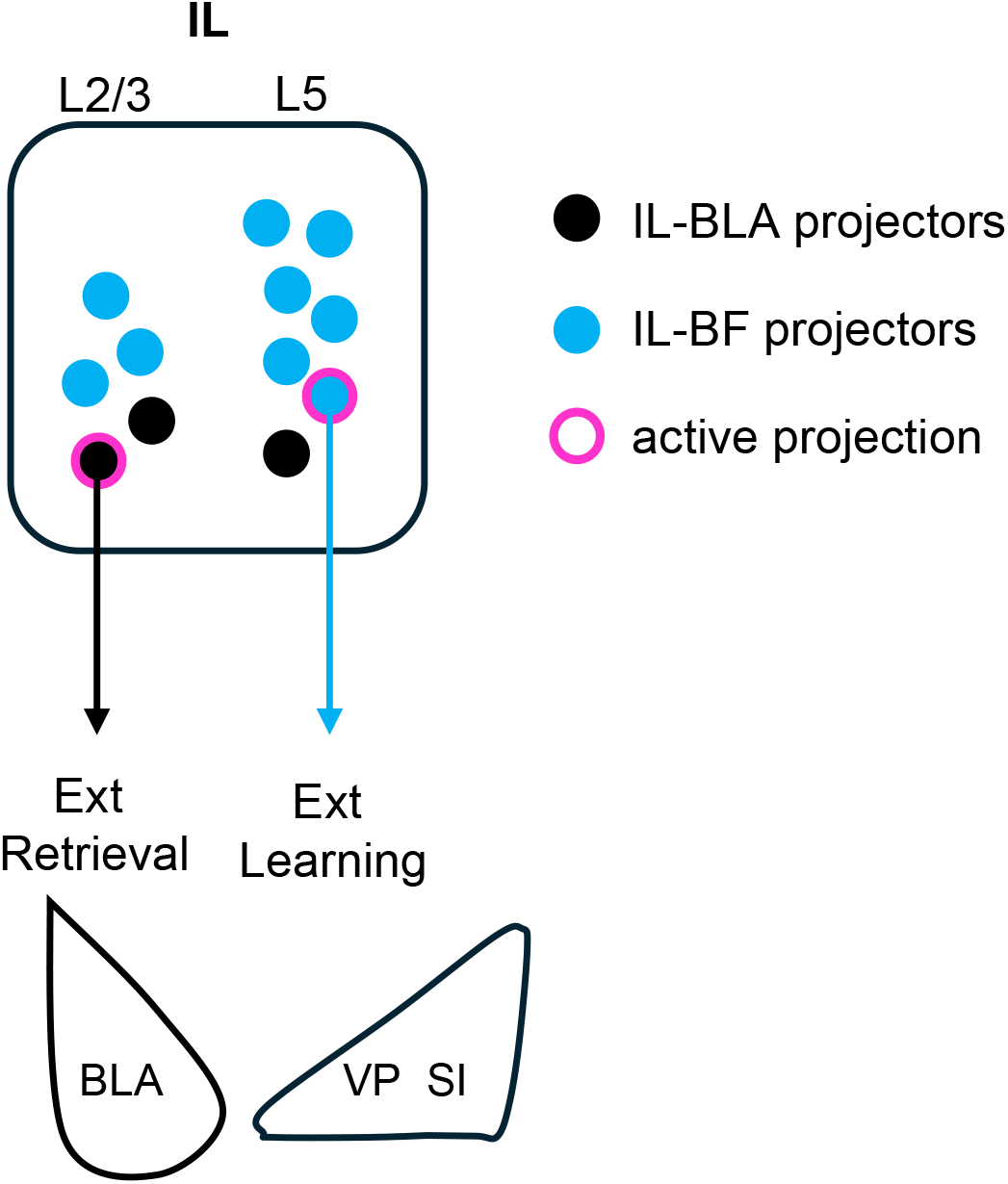
Summary model. Neuroanatomical distribution of IL projectors to the BLA (black circles) and BF (blue circles) across layers, with L5 IL-BF projectors becoming more active (pink rim) during extinction learning, and L2/3 IL-BLA projectors becoming more active (pink rim) during extinction retrieval.

Furthermore, we demonstrate that inhibiting the IL-to-BF output during extinction learning slows extinction learning but does not affect extinction memory retrieval. Taken together, we show, for the first time, that as extinction progresses, IL outputs to the BF become more active, and IL communication with the BF partakes in suppressing fear expression during learning.

The IL projects to targets that are cortical (e.g. insula), subcortical (e.g. BF, BLA, the paraventricular nucleus of the thalamus [PVT], hypothalamus), and in the brainstem (e.g. the periaqueductal grey, nucleus of the solitary tract), which collectively influence cardiovascular, motor, and cognitive responses (Room et al. 1985, Hurley et al. 1991, Vertes 2004). This intriguing map of IL targets lead us to interrogate whether IL projections to the BF, a major source of cholinergic innervation throughout the central nervous system, has a role to play in extinction. Interestingly, optogenetic inactivation of the IL during extinction retrieval was reported not to impair extinction memory (Do-Monte et al. 2015), suggesting that the IL facilitates extinction retrieval via its target regions. The role of the IL-BLA pathway in mediating extinction retrieval has received the most attention due to the critical role of the BLA in fear processing (Quirk et al. 2003, Amano et al. 2010, Cho et al. 2013, Strobel et al. 2015). Indeed, previous work has shown that inhibition of the IL-BLA pathway during extinction learning doesn’t impair within-session extinction, but instead dampens extinction memory (Bukalo et al. 2015, Bloodgood et al. 2018). Likewise, recent work has uncovered the PVT as another important target of the IL for mediating extinction retrieval (Tao et al. 2021). Notably, *in vitro* recordings of cells in the IL, including in IL-BLA projectors, show that they increase in excitability after extinction (Santini et al. 2008, Cruz et al. 2014, Bloodgood et al. 2018), suggesting cellular and molecular consolidation processes occur in IL-BLA projectors post-extinction learning (Santini et al. 2004, Burgos-Robles et al. 2007), which could then alter post-synaptic activity at target sites that handle a variety of extinction-related behavioral components.

Less is known about the IL pathway to the BF and its role in extinction. Our comparison of cFos activity in the IL-BLA vs IL-BF pathways shows that the IL-BLA L2/3 pathway is upregulated during extinction retrieval, as would be expected from previous work (Bukalo et al. 2015, Bloodgood et al. 2018). Conversely, the IL-BF L5 pathway is upregulated during extinction learning. We used *in vitro* recordings for a more granular temporal analysis of IL-BF projector excitability. These recordings showed that IL-BF projectors became more excitable as extinction learning progressed. Notably, excitability increased with no change in the resting membrane potential of IL-BF projectors, suggesting that their increased excitability is due to network effects rather than intrinsic changes. Increased network activation could come from several inputs to the IL, including the ventral hippocampus (Kim and Cho 2017, Brockway et al. 2023), the PVT (Russo and Parsons 2022), and the prelimbic cortex (Marek et al. 2018, Watanabe et al. 2021). Thus emerges a picture of brain-wide communication during extinction, whereby IL afferents shape its activity during learning, and a variety of its efferents drive memory retrieval. However, this is the first demonstration of a pathway whereby the IL can downregulate fear expression during extinction learning.

The BF is one of the main sources of brain-wide cholinergic innervation, targeting regions in the fear processing circuit such as the amygdala, the hippocampus, as well as the auditory and prefrontal cortices (Zaborszky et al. 2015, Gielow and Zaborszky 2017). From a cognitive perspective, BF cholinergic signaling drives attentional processing of cues (Parikh et al. 2007, Gritton et al. 2016), which in extinction, include the CS and/or the context.

Interestingly, lesions of the cholinergic BF were previously shown to impair cued within-session extinction (Knox 2016, Knox and Keller 2016) and inhibition of cholinergic signaling impaired contextual extinction learning (Zelikowsky et al. 2013), indicating that BF cholinergic activity is necessary for extinction acquisition. However, BF cholinergic signaling is also upregulated during fear learning. For example, there is cholinergic release in the BLA during fear learning (Rajebhosale et al. 2024), which can counteract fear extinction (Jiang et al. 2016), increasing the BLA signal-to-noise patterns of neural firing (Unal et al. 2015, Knox 2016), and entraining BLA theta oscillations that are associated with threat (Aitta-Aho et al. 2018, Bratsch-Prince et al. 2024, Cattani et al. 2024). Thus, given the importance of cholinergic signaling for both fear and extinction, it’s likely that cholinergic tone helps modulate attention to the CS during multiple kinds of learning (Likhtik and Johansen 2019). For example, it was shown that cholinergic signaling in the BLA is important for forming reward associations with a CS (Crouse et al. 2020). However, too much cholinergic signaling is associated with anxiety and depression, such that it’s necessary to strike a homeostatic balance that facilitates focus during learning but prevents overactivation (Vythilingam et al. 2007, Mineur and Picciotto 2021). Our finding that inhibiting IL projections to the BF kept defensive freezing high during extinction learning suggests that this pathway could serve as an important means for down-regulating cholinergic BF output as extinction learning progresses.

Extinction is a form of prediction-error based learning, a model of which was first formalized by Rescorla and Wagner (1972). During extinction, the omission of the US violates the predicted CS-US association that was established during fear conditioning, facilitating new learning about the CS (reviewed in (Bouton et al. 2021)). Notably, prediction-error models highlight the importance of attention to the CS for learning the changing CS-US association (Mackintosh 1975, Pearce and Hall 1980, Dunsmoor et al. 2015), which is reflected in increased attentional network activity in humans during extinction learning (Wen et al. 2021). Given that acetylcholine signaling increases when a known CS is unreliable (Yu and Dayan 2005), we would expect the highest cholinergic activity at the beginning of extinction, when the previously established CS-US contingency is first violated. The CS is therefore uncertain, and the prediction error is high. Notably, the cholinergic BF sends an important projection to the IL (Henny and Jones 2008, Bloem et al. 2014, Zaborszky et al. 2015), which regulates extinction consolidation via cholinergic receptors in the prefrontal cortex (Santini et al. 2012, Wilson and Fadel 2017). Thus exists a neuroanatomical loop between the cholinergic BF and the IL, which could support attention related processing from the bottom-up and from the top-down (Sarter et al. 2005). Interestingly, these two streams are likely to interact with each other, affecting cholinergic modulatory strength (Sarter et al. 2005). In such a loop, BF cholinergic communication with regions such as the mPFC and BLA could be part of bottom-up signaling about the unreliable CS at the beginning of learning. Then, the IL-to-BF signaling could partake in the top-down process that helps downmodulate cholinergic activity and diminish within-session fear expression. Notably, previous work has shown evidence for prefrontal interactions with attention network in emotion regulation (Sharpe and Killcross 2014, Sharpe and Killcross 2015, Wen et al. 2021), suggesting that this an important loop for cognitive control during learning.

During extinction retrieval, the IL-to-BLA pathway suppresses fear by driving heterosynaptic inhibition of CS-driven activity in the BLA, and diminishing amygdala output (Royer and Pare 2002, Quirk et al. 2003, Likhtik et al. 2008, Amano et al. 2010, Amir et al. 2011, Cho et al. 2013, Strobel et al. 2015). This suppression involves both excitatory and inhibitory networks within the BLA’s microcircuitry, highlighting a vital physiological mechanism for controlling fear during extinction retrieval. A non-overlapping, but similarly intricate set of IL interactions with the BF microcircuit, which contains cholinergic, inhibitory, and glutamatergic cells (Gritti et al. 2006), could be occurring during extinction acquisition. Notably, evidence suggests that electrical stimulation of the IL drives fast-spiking neurons in the BF while decreasing activity of slower-spiking cells, which could reflect cholinergic firing (Gyengesi et al. 2008). Understanding how the IL impacts the BF microcircuit may offer important insights into how the IL mediates BF activity, attention, and fear expression during extinction learning.

## Acknowledgements

This work was supported by a National Institutes of Health research program grant (MH118441). The authors would like to thank Dr. Sarah Mennenga for help with statistical modeling, Dr. Andrew Delamater for helpful discussions of the manuscript, and Itzick Nahmoud for help with programming of the behavioral equipment.

